# The mitotic kinesin-14 KlpA contains a context-dependent directionality switch

**DOI:** 10.1101/058602

**Authors:** Andrew R. Popchock, Kuo-Fu Tseng, Pan Wang, P. Andrew Karplus, Xin Xiang, Weihong Qiu

**Author notes:** These authors contributed equally (Andrew R. Popchock, Kuo-Fu Tseng).

## Abstract

Kinesins are microtubule-based motor proteins that convert chemical energy from ATP hydrolysis into mechanical work for a variety of essential intracellular processes. Kinesin-14s (i.e. kinesins with a C-terminal motor domain) are commonly considered to be nonprocessive minus end-directed motors that mainly function for mitotic spindle assembly and maintenance. Here, we show that KlpA – a mitotic kinesin-14 motor from the filamentous fungus *Aspergillus nidulans* – contains a context-dependent directionality switch. KlpA exhibits canonical minus end-directed motility inside microtubule bundles, but on individual microtubules it unexpectedly moves processively toward the plus ends. Removal of the N-terminal nonmotor microtubule-binding domain renders KlpA diffusive on individual microtubules but does not abolish its minus end-directed motility to collectively glide microtubules, suggesting that the nonmotor microtubule-binding domain likely acts as a switch for controlling the direction of KlpA motility. Collectively, these findings provide important insights into the mechanism and regulation of KlpA functions inside the mitotic spindle.

## Introduction

The mitotic spindle is a microtubule-based bipolar machine in eukaryotes that separates duplicated chromosomes to ensure that daughter cells each receive proper genetic material during cell division. Several different kinesin motor proteins are orchestrated inside the mitotic spindle for its assembly and maintenance^1,2^. Of all mitotic kinesins, kinesin-14s are commonly considered to be nonprocessive minus end-directed microtubule motors^3–12^. While mitotic kinesin-14s are nonessential for normal cells, loss of the kinesin-14 Pkl1 in fission yeast *Schizosaccharomyces pombe* has been shown to cause erroneous chromosome segregation^13^. In cancer cells, the human kinesin-14 HSET/KIFC1 is needed for clustering multiple centrosomes, a process crucial for cancer cell proliferation and survival^14^.

KlpA is a mitotic kinesin-14 from the filamentous fungus *Aspergillus nidulans*^15^. It is worth noting that *A. nidulans* is also the model organism for the discovery of BimC, the founding member of mitotic kinesin-5s^16^. Like mitotic kinesin-14s in other eukaryotic cells^11,17,18^, KlpA counteracts the function of BimC^15^. Similar to the fission yeast kinesin-14 Pkl1^19^, KlpA is nonessential in wildtype cells but its loss becomes synthetically lethal with gamma tubulin mutations^20^. KlpA is an attractive model protein for dissecting the mechanism and function of kinesin-14s, as its loss-of-function mutations can be conveniently isolated as suppressors of the bimC4 mutation^21^. However, compared with other mitotic kinesin-14s such as Ncd from *Drosophila melanogaster* and Kar3 from *Saccharomyces cerevisiae,* KlpA is much less well studied.

In this study, we report our *in vitro* characterization of KlpA motility using total internal reflection fluorescence (TIRF) microscopy. KlpA unexpectedly moves processively toward the plus ends on individual microtubules as a single homodimer and switches to the canonical minus end-directed motility inside microtubule bundles. Thus, KlpA is a context-dependent bidirectional kinesin-14, making it distinct from all other kinesin-14s that have been examined to date. Furthermore, our results suggest that KlpA contains an N-terminal nonmotor microtubule-binding domain that not only enables the motor for plus end-directed processive motility but also acts a switch for controlling its directionality in different cellular contexts. These findings shed new light on KlpA motor mechanism and provide a molecular view of how KlpA may be regulated for mitotic spindle assembly and maintenance.

## Results

### KlpA glides microtubules with canonical minus end-directed motility

We set out to determine the directionality of KlpA *in vitro* using TIRF microscopy. To that end, we purified the recombinant full-length KlpA tagged with an N-terminal green fluorescent protein (GFP-KlpA, Fig. 1a, b). Since KlpA substitutes for Kar3 in *S. cerevisiae*^15^ and Kar3 forms a heterodimer with the nonmotor proteins Cik1 or Vik1^22^, we performed two different assays – hydrodynamic analysis and single-molecule photobleaching – to determine the oligomerization status of KlpA. The hydrodynamic analysis yielded a molecular weight that is close to the theoretical value of a GFP-KlpA homodimer (Supplementary Fig. 1a, b). The photobleaching assay showed that the GFP fluorescence of GFP-KlpA was predominantly photobleached in a single step or two steps (Supplementary Fig. 1c, d), similar to other dimeric kinesins^23^. Thus, unlike *S. cerevisiae* kinesin-14 Kar3^22^ but similar to other kinesin-14s such as *D. melanogaster* Ncd^24^ and *S. pombe* Klp2^25^, KlpA forms a homodimer.

**Fig 1:**
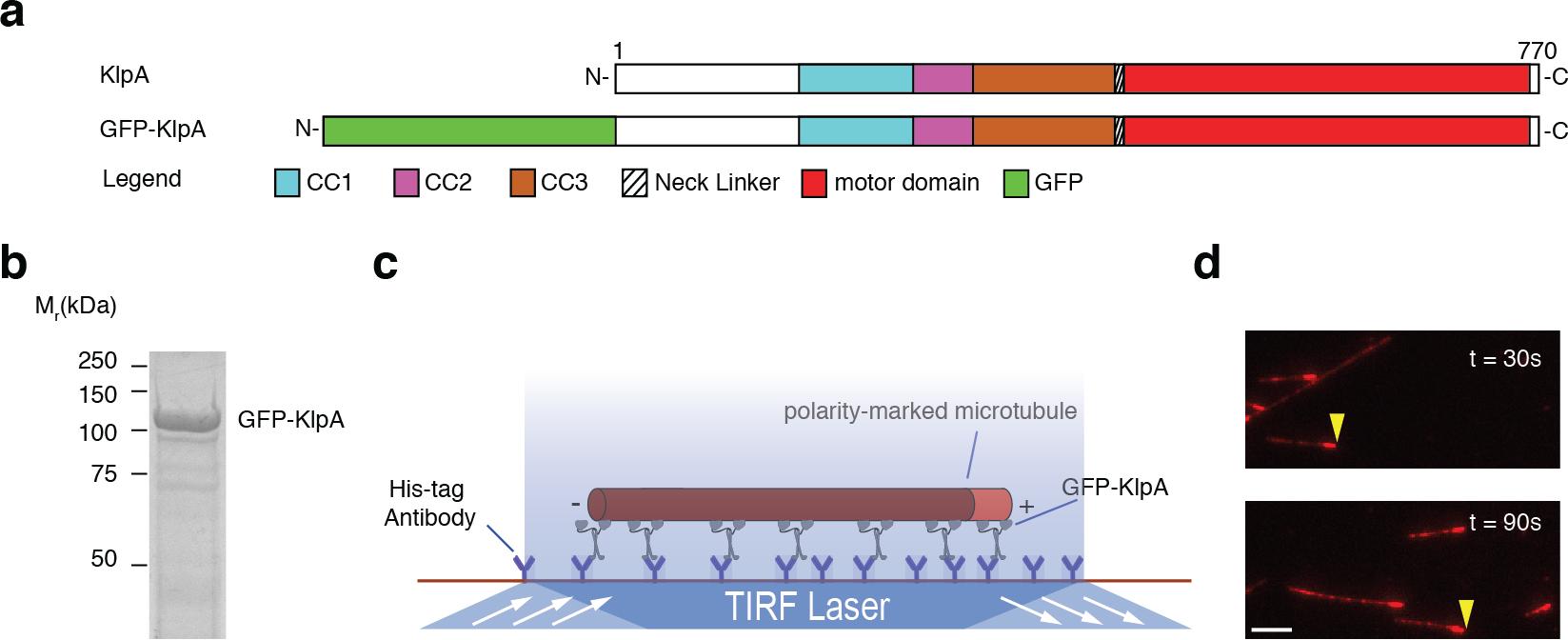
Surface-immobilized KlpA molecules exhibit minus end-directed motility to glide microtubules. **a**, Schematic diagrams of the full-length KlpA and the recombinant GFP-KlpA. The full-length KlpA consists of three consecutive coiled coils (CC1, aa 153-249; CC2, aa 250-297; and CC3, aa 298-416), a neck linker (aa 417-421), and a catalytic microtubule-binding motor domain (aa 422-756). GFP-KlpA contains an N-terminal polyhistidine-tag (not shown). **b**, Coomassie-stained SDS-polyacrylamide gel electrophoresis (SDS-PAGE) of purified recombinant GFP-KlpA. **c**, Schematic diagram of the microtubule-gliding assay. Movement of microtubules driven by surface-immobilized GFP-KlpA molecules was visualized by total internal reflection fluorescence (TIRF) microscopy. Microtubules were fluorescently labeled with tetramethylrhodamine (TMR), and polarity-marked with a dim minus end and a bright plus end^48^. **d**, Representative TIRF microscopy images of polarity-marked microtubules gliding with the bright plus ends leading (yellow arrowheads). Scale bar: 5 ¼m.

We next performed a microtubule-gliding assay to determine the directionality of KlpA (Fig. 1c). Briefly, GFP-KlpA molecules were immobilized on the coverslip via an N-terminal polyhistidine-tag, and KlpA directionality was deduced from the motion of polarity-marked microtubules. The assay showed that GFP-KlpA caused polarity-marked microtubules to move with the bright plus ends leading (Fig. 1d and Supplementary Video 1). In a control experiment using the plus end-directed human conventional kinesin hKHC^26^, microtubules were driven to move with the bright plus ends trailing (Supplementary Fig. 2 and Supplementary Video 2). Taken together, these results demonstrate that KlpA, anchored on the surface via its N-terminus, is a minus end-directed motor protein, in agreement with a previous study using KlpA from clarified bacterial lysates^20^.

### Single KlpA molecules move processively toward the plus ends on individual microtubules

We wanted to determine whether KlpA is a typical kinesin-14 that lacks the ability to move processively on individual microtubules as a single homodimer. To address this, we performed an *in vitro* motility assay to visualize the movement of KlpA molecules on surface-immobilized polarity-marked microtubules (Fig. 2a). The assay was first performed at relatively high input levels of GFP-KlpA (≥ 4.5 nM). Contrary to the notion of kinesin-14s as minus end-directed motors, GFP-KlpA molecules unexpectedly formed a steady flux to accumulate at the microtubule plus ends (yellow arrow, Fig. 2b and Supplementary Video 3). Occasionally, there were GFP-KlpA particles moving toward the microtubule minus ends (white arrow, Fig. 2b), but these minus end-directed particles were significantly brighter than the ones moving toward the plus ends, implying that they were aggregates rather than simple homodimers. Since GFP-KlpA appeared to move processively toward the microtubule plus ends (Fig. 2b), we repeated the *in vitro* motility assay at lower protein input levels (≤ 0.2 nM) so that the motile behavior of individual GFP-KlpA molecules could be distinguished. The assay showed that individual GFP-KlpA molecules moved preferentially toward the microtubule plus ends in a processive manner (Fig. 2c and Supplementary Video 4) with a mean velocity of ~320 ± 90 nm/s (mean ± s.d., n = 249, Fig. 2d) and a characteristic run-length of 8.8 ± 0.2 ¼m (mean ± s.e., n = 249, Fig. 2e). This run-length likely was an underestimate, as most KlpA molecules reached the microtubule plus ends. Together, these results demonstrate that KlpA, in direct contrast to all other kinesin-14s examined to date, is a processive plus end-directed kinesin.

**Fig 2:**
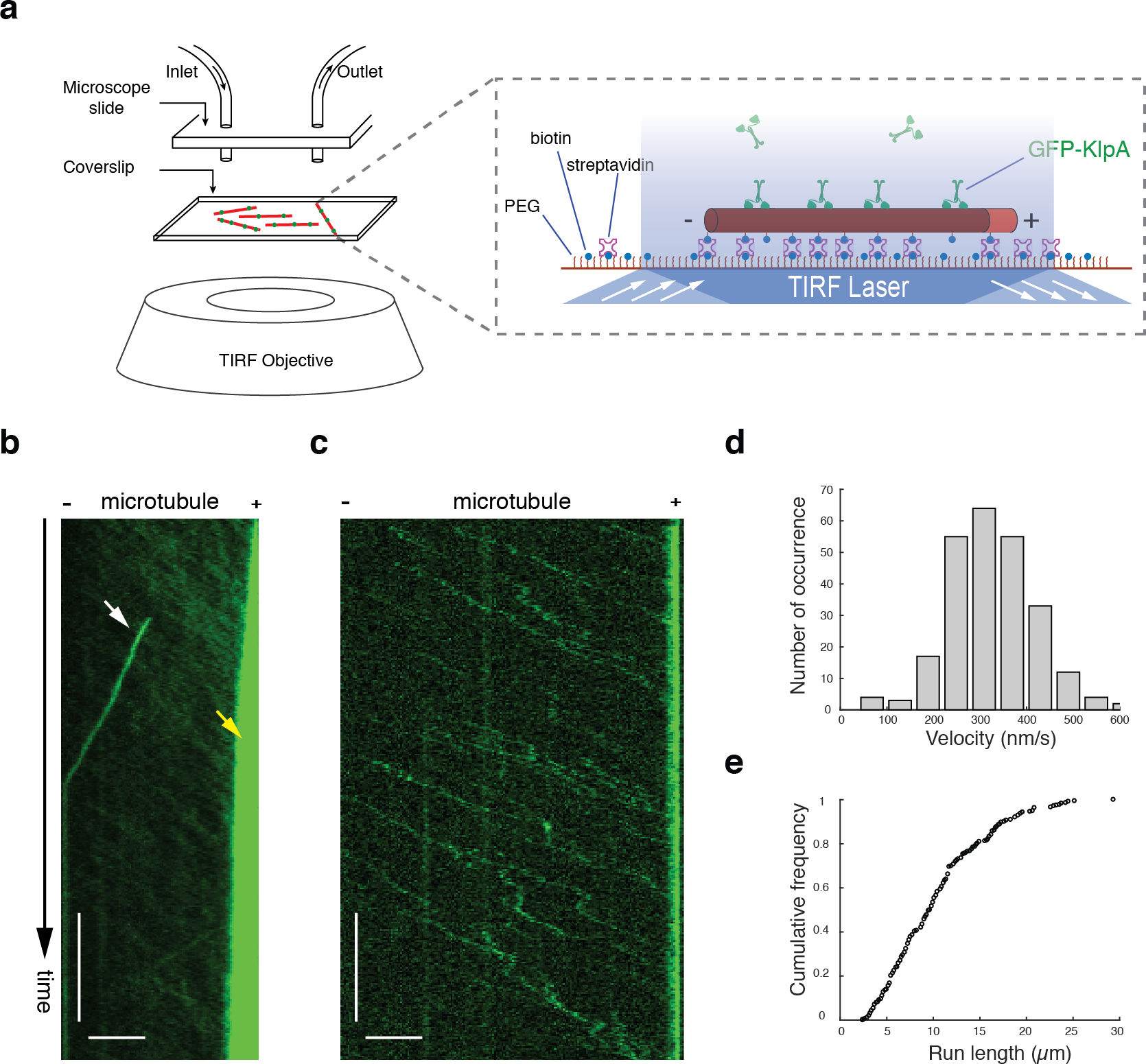
Single KlpA molecules move processively toward the plus ends on individual microtubules. **a**, Schematic diagram of the *in vitro* KlpA motility assay. Microtubules were fluorescently labeled with Hilyte 647, and polarity-marked with a dim minus end and a bright plus end^48^. **b**, Example kymograph showing GFP-KlpA molecules, at relatively high protein input levels, form a plus end-directed flux and accumulate there on individual microtubules. Yellow arrow indicates GFP-KlpA accumulation at the microtubule plus end, and white arrow indicates minus-end-directed movement of a GFP-KlpA aggregate. **c**, Example kymograph showing that single GFP-KlpA molecules move preferentially toward the plus end on individual microtubules in a processive manner. **d**, Histogram of the velocity of GFP-KlpA. **e**, Cumulative frequency of the run-length of GFP-KlpA. Scale bars: 1 minute (vertical) and 5 ¼m (horizontal).

### An N-terminal nonmotor microtubule-binding domain in KlpA is required for its plus end-directed processive motility

Like other kinesin-14s such as Klp2 in *S. pombe* and Ncd in *D. melanogaster*^24,25^, KlpA was also able to slide antiparallel microtubules and to statically crosslink parallel microtubules via a nonmotor microtubule-binding domain (MTBD) at the N-terminus (Supplementary Fig. 3a-g, and Supplementary Video 5 and 6). As several other kinesins are known to rely on nonmotor MTBDs to either achieve processive motility^22^ or enhance processivity^27,28^, we sought to determine whether the N-terminal nonmotor MTBD of KlpA plays a similar role to its unexpected plus end-directed processive motility. To do this, we purified GFP-KlpA-Δtail (Fig. 3a), a truncated construct lacking the N-terminal nonmotor MTBD, for *in vitro* motility experiments. Like GFP-KlpA, GFP-KlpA-Δtail formed a homodimer (Supplementary Fig. 4) and exhibited minus end-directed motility in the microtubule-gliding assay (Fig. 3b and Supplementary Video 7). This latter observation implies that the motor core of KlpA without the N-terminal nonmotor MTBD is inherently minus end-directed, as would be expected based on its highly conserved neck linker ^26,29–31^. However, the *in vitro* motility assay showed that GFP-KlpA-Δtail did not form a steady flux toward either end of the microtubule, nor did it accumulate at the microtubule ends (Fig. 3c and Supplementary Video 8). Although some occasional brighter and presumably aggregated particles moved processively toward the microtubule minus ends (white arrow, Fig. 3c and Supplementary Video 8), individual GFP-KlpA-Δtail molecules interacted with the microtubules in a diffusive manner with no obvious directional preference. Thus, besides allowing for microtubule sliding and crosslinking, the N-terminal nonmotor MTBD has an additional novel functionality of enabling KlpA to move on individual microtubules toward the plus ends in a processive manner.

**Fig 3:**
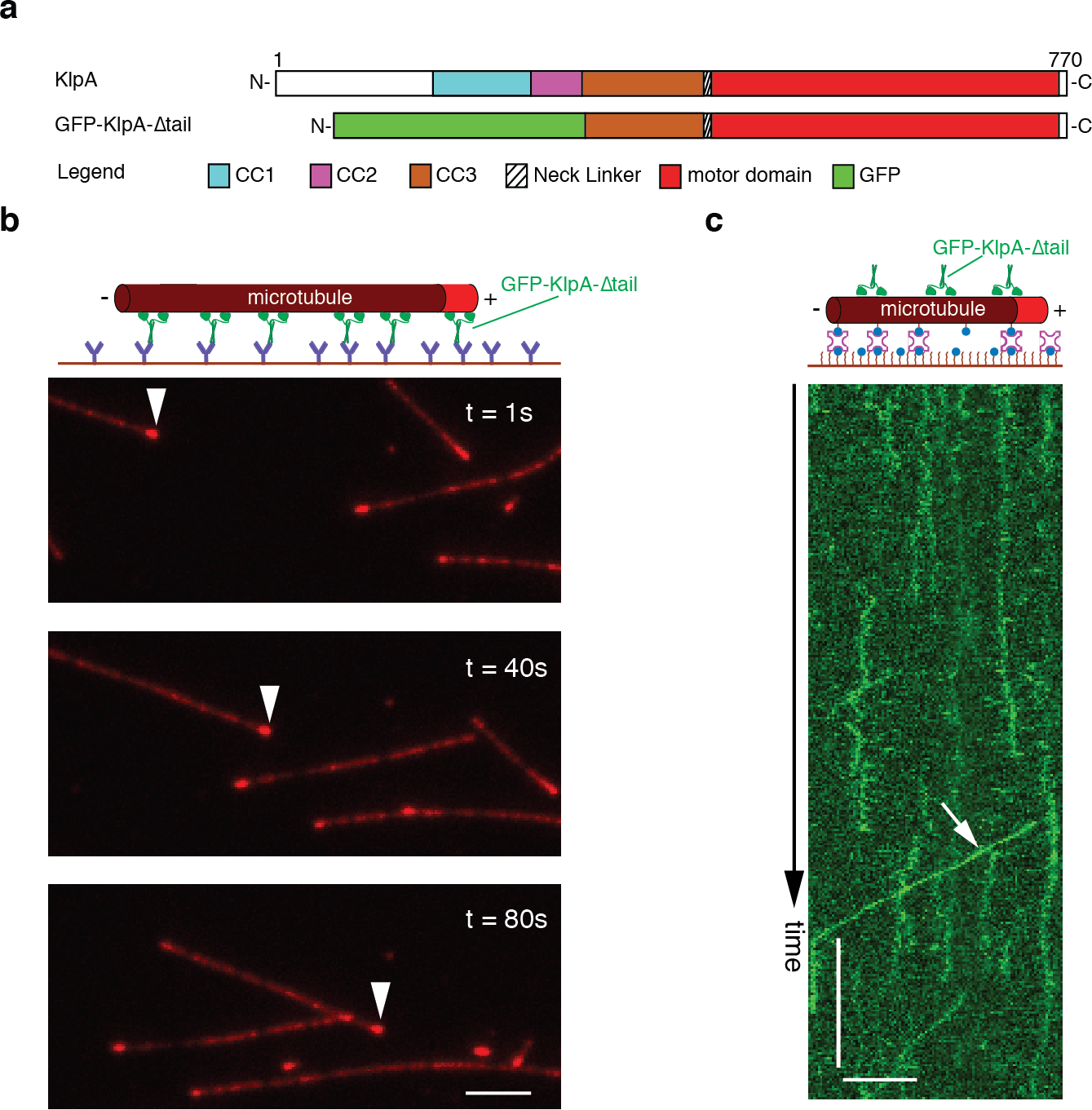
The N-terminal nonmotor MTBD of KlpA is required for its plus end-directed motility on individual microtubules. **a**, Schematic diagrams of the full-length KlpA and the recombinant GFP-KlpA-Δtail. GFP-KlpA-Δtail contains a polyhistidine-tag (not shown) and a GFP at the N-terminus, and residues 303-770 of KlpA. **b**, Representative TIRF microscopy images of GFP-KlpA-Δtail driving polarity-marked microtubules to glide with the bright plus ends leading (white arrowheads). **c**, Example kymograph showing that GFP-KlpA-Δtail lacks plus end-directed motility and mostly diffuse on the microtubules. White arrow indicates minus-end-directed movement of a rare GFP-KlpA-Δtail aggregate. Scale bars: 1 minute (vertical) and 5 ¼m (horizontal).

### KlpA exhibits opposite directional preference inside and outside microtubule overlaps

From the opposite directional preference exhibited by GFP-KlpA in the ensemble microtubule assay (Fig. 1d and Supplementary Fig. 3c) and the single-molecule motility experiments (Fig. 2b, c), we inferred that KlpA contains a context-dependent mechanism to switch directions on the microtubule^32^. We thus directly compared the motility of GFP-KlpA inside and outside the microtubule overlap on the same track microtubule using a microtubule-transport assay (Fig. 4a), as has been done previously for *S. cerevisiae* kinesin-5 Cin8^32^. Briefly, in this assay the track (blue) and cargo (red) microtubules were both polarity-marked but labeled with different dyes; track microtubules were first immobilized on a coverslip inside the motility chamber and bound with purified GFP-KlpA molecules; and cargo microtubules were added into the chamber before three-color time-lapse imaging was acquired to simultaneously visualize the motility of GFP-KlpA molecules and cargo microtubules on the same track microtubules. Like KlpA, GFP-KlpA was also able to slide antiparallel microtubules relative to each other (Fig. 4b) and to statically crosslink parallel microtubules (Fig. 4c). In both scenarios, when outside the microtubule overlap regions, GFP-KlpA molecules showed a plus end-directed flux and accumulated at the plus end on the track microtubule (yellow arrow, Fig. 4b, c and Supplementary Video 9 and 10). This matches the behavior of GFP-KlpA on individual microtubules (Fig. 2b). In contrast, inside the antiparallel microtubule overlap regions, GFP-KlpA molecules carried the cargo microtubule toward the minus end of the track microtubule (white arrow, Fig. 4b and Supplementary Video 9). In the parallel orientation, the cargo microtubule remained stationary on the track microtubule, but GFP-KlpA molecules moved preferentially toward and gradually accumulated at the minus end inside the parallel microtubule overlap (white arrow, Fig. 4c and Supplementary Video 10). This is similar to the observation that Ncd preferentially accumulates at the minus ends between statically crosslinked parallel microtubules^24^. Collectively, these results demonstrate that KlpA can, depending on context, display opposite directional preferences on the same microtubule: it is plus end-directed outside the microtubule overlap regions and minus end-directed inside the microtubule overlap regions regardless the relative microtubule polarity.

**Fig 4:**
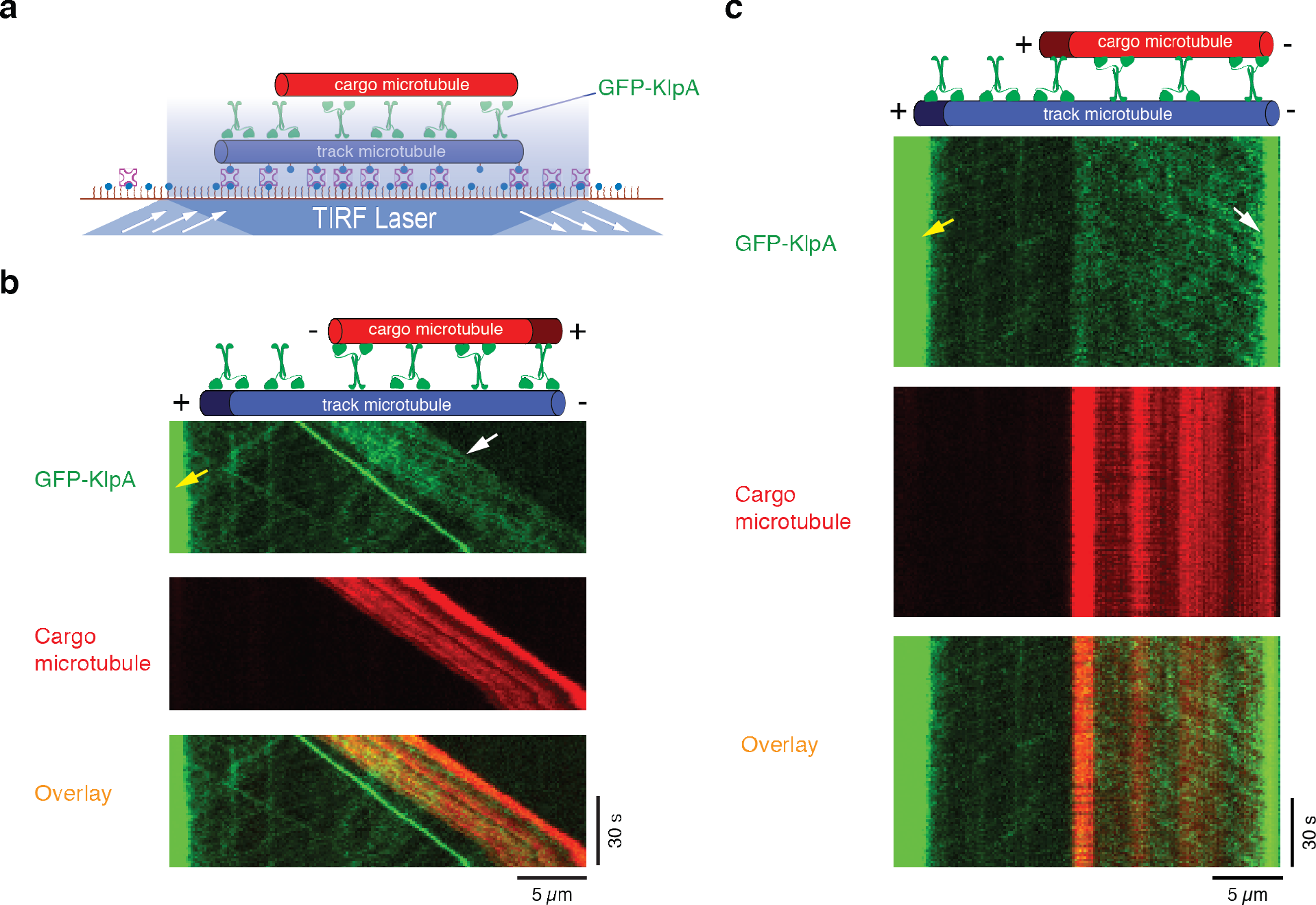
KlpA exhibits opposite directional preference inside and outside the microtubule overlaps. **a**, Schematic diagram of the microtubule-transport assay showing that KlpA contains context-dependent opposite directional preference. Track and cargo microtubules were fluorescently labeled with Hilyte 647 and TMR respectively and polarity-marked with a dim minus end and a bright plus end^48^. **b**, Example kymographs of GFP-KlpA motility inside and outside the antiparallel microtubule overlap. Yellow arrow indicates GFP-KlpA accumulation at the microtubule plus end outside the antiparallel microtubule overlap. White arrow indicates minus-end-directed movement of GFP-KlpA inside the antiparallel microtubule overlap **c**, Example kymographs of GFP-KlpA motility inside and outside the parallel microtubule overlap. Yellow arrow indicates GFP-KlpA accumulation at the microtubule plus end outside the parallel microtubule overlap. White arrow indicates GFP-KlpA accumulation at the microtubule minus end inside the parallel microtubule overlap.

## Discussion

Kinesin-14 has been an intriguing kinesin subfamily since the discovery of its founding member Ncd^33,34^, because all kinesin-14s studied to date are exclusively minus end-directed in the microtubule-gliding experiments^23,25,33–39^. With the lone exception of Kar3, no other kinesin-14 has been shown to be able to generate processive motility directly on the surface of individual microtubules as a single homodimer. In *vitro*, it has been shown that Kar3 generates processive minus end-directed motility on individual microtubules by forming a heterodimer with its associated light chains Vik1 or Cik1^22,40^. By revealing KlpA as a kinesin-14 that demonstrates both processive plus end-directed motility on individual microtubules and context-dependent directional switching, our study further expands the diversity of kinesin-14s.

How does KlpA achieve the observed context-dependent directional switching? Our results show that while the full-length KlpA clearly moves processively toward the plus ends on individual microtubules (Fig. 2b, c), a truncated KlpA lacking the N-terminal nonmotor MTBD is unable to produce processive motility (Fig. 3c) but does retain the ability to glide microtubules with minus end-directed motility (Fig. 3b). There are several important implications from these observations. First, the motor core of KlpA without the nonmotor MTBD is inherently minus end-directed, which is in agreement with the notion that all kinesin-14s share a highly conserved neck linker that serves as the minus end directionality determinant^26,29–31^. Second, the nonmotor MTBD is required for plus end-directed KlpA motility on individual microtubules. We suggest that the nonmotor MTBD is a *de facto* switch for controlling the direction of KlpA motility: KlpA is plus end-directed kinesin-14 motor when the switch-like nonmotor MTBD and the motor domain both bind to the same microtubule, and it reverses to become a nonprocessive minus end-directed motor when the switch-like nonmotor MTBD is detached from the microtubule to which its motor domain binds. This could explain the minus end-directed motility of KlpA anchored on the coverslip via the N-terminus (Fig. 1d) or inside microtubule bundles (Fig. 4b, c), because in both cases the switch-like nonmotor MTBD is in effect detached from the microtubule to which its motor domain binds. Future studies will need to determine the structural basis of how the nonmotor MTBD enables KlpA to move with plus end-directed processive motility. We speculate that positioning of the nonmotor MTBD relative to the motor domain on the microtubule may favor KlpA to search the next binding site between steps toward the microtubule plus ends.

Our findings provide a molecular view for how KlpA motility may be regulated inside the mitotic spindle (Fig. 5). While other mitotic kinesin-14s appear to depend on partner proteins to localize to the spindle midzone for antagonizing the action of kinesin-5s^10,37,41,42^, KlpA can in principle autonomously localize to the spindle midzone via its inherent plus end-directed motility by having both the nonmotor MTBD and the motor domain on the same microtubule (Fig. 5a). Inside the antiparallel microtubule overlaps at the spindle midzone (Fig. 5b) or the parallel microtubule overlaps near the spindle poles (Fig. 5c), KlpA switches to become minus end-directed as the switch-like nonmotor MTBD and the motor domain bind to two different microtubules. This apparent directional plasticity suggests that other proteins could exist to regulate KlpA motility via intermolecular interactions that interfere with the binding of the switch-like nonmotor MTBD to microtubules. A recent study shows that Pkl1 – a mitotic kinesin-14 from the fission yeast - forms a complex with Msd1 and Wdr8 for translocating to and anchoring at the spindle poles^43^. The homologs of both Msd1 and Wdr8 are also present in *A. nidulans*^44^. Thus, it is plausible that binding of Msd1 and Wdr8-like proteins to KlpA could dislodge its N-terminal nonmotor MTBD from the surface of microtubules to activate the kinesin for minus end-directed motility both on individual microtubules (Fig. 5d) and at the spindle poles (Fig. 5e).

**Fig 5:**
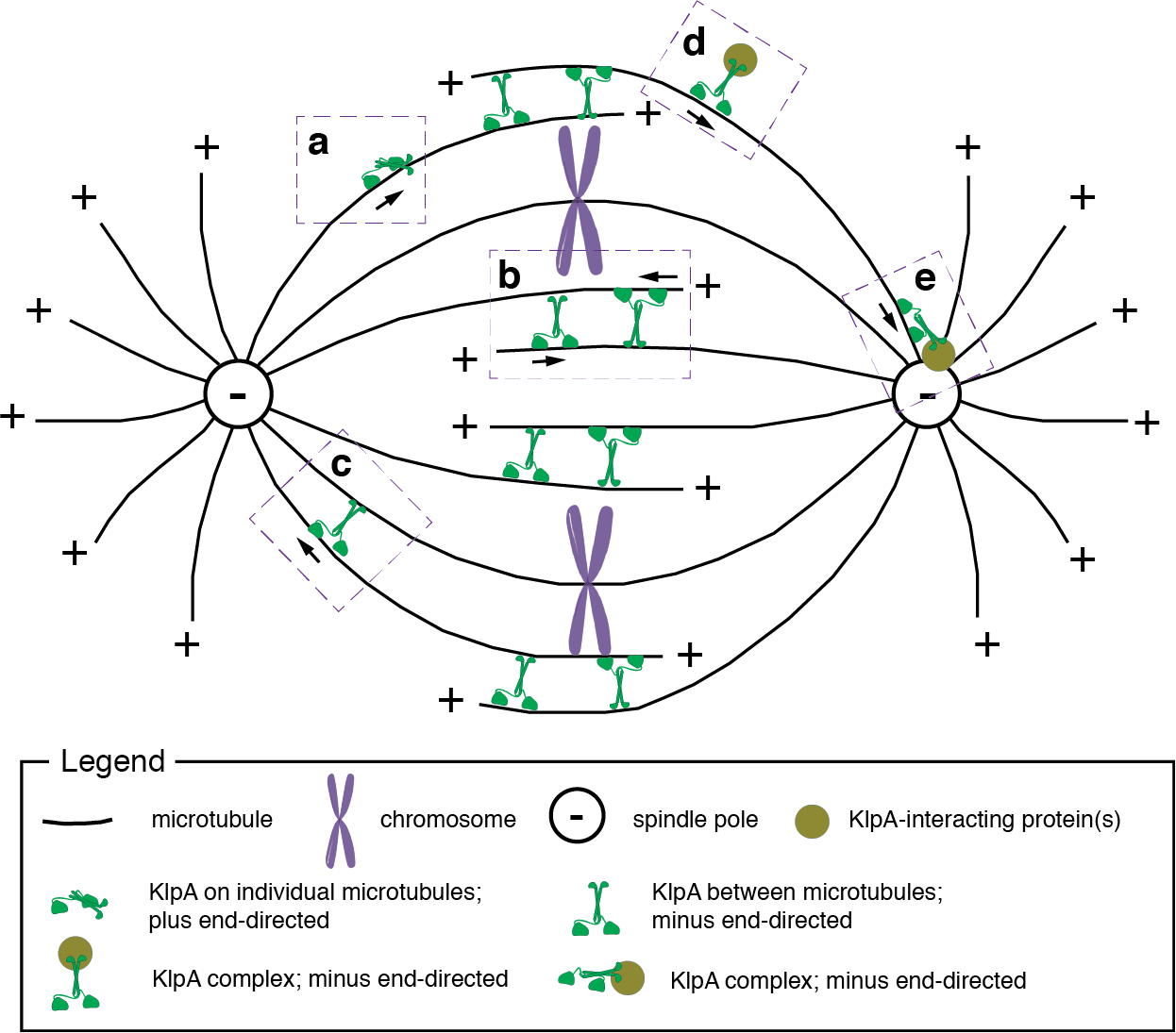
Schematic model illustrating the context-dependent directional switching of KlpA inside the mitotic spindle. **(a)**, KlpA moves preferentially toward the plus end in a processive manner on individual microtubules. **(b-e)**, KlpA is minus end-directed between antiparallel microtubule overlap (**b**), between parallel microtubule overlap (**c**), in complex with its putative cargo protein(s) on individual microtubules (**d**), and anchored at the spindle pole via its cargo protein(s) (**e**).

Several mitotic kinesin-5s were recently shown to be context-dependent bidirectional motor proteins^32,45–47^, suggesting that context-dependent directional switching likely is evolutionarily conserved among kinesin-5s. Our current work on KlpA provides the first evidence to suggest that context-dependent directional switching could also exist among some, if not all, mitotic kinesin-14s. The mechanism and regulation of bidirectional mitotic kinesins will be an important subject for future studies.

## Methods

Detailed methods are described in Supplementary Information.

## Acknowledgements

We thank Drs. C. Mathews (Oregon State University), X. Su (UCSF) and B. Liu (UC Davis) for critical reading of the manuscript, and Mr. Chun Liu (Pearl River Fisheries Research Institute, China) for initial plasmid construction.

## Author Contributions

W.Q. conceived, designed and supervised the study; P.A.K. and X.X. provided conceptual suggestions; A.R.P. and K.-F.T. performed the experiments; K.-F.T. and P.W. contributed all KlpA constructs. All authors participated in discussing the results. W.Q. wrote the manuscript with input from all authors.

## Author Information

The authors declare no competing financial interests. Correspondence and requests for materials should be addressed to.

## References

1. Wordeman, L. How kinesin motor proteins drive mitotic spindle function: Lessons from molecular assays. Semin. Cell Dev. Biol. 21, 260–268 (2010).

2. Winey, M. & Bloom, K. Mitotic spindle form and function. Genetics 190, 1197–1224 (2012).

3. Marcus, A. I., Li, W., Ma, H. & Cyr, R. J. A kinesin mutant with an atypical bipolar spindle undergoes normal mitosis. Mol. Biol. Cell 14, 1717–1726 (2003).

4. Ambrose, J. C. & Cyr, R. The kinesin ATK5 functions in early spindle assembly in Arabidopsis. Plant Cell 19, 226–236 (2007).

5. Endow, S. A. & Higuchi, H. A mutant of the motor protein kinesin that moves in both directions on microtubules. Nature 406, 913–916 (2000).

6. Endow, S. A. & Komma, D. J. Centrosome and spindle function of the Drosophila Ncd microtubule motor visualized in live embryos using Ncd-GFP fusion proteins. J. Cell Sci. 109, 2429–2442 (1996).

7. Hatsumi, M. & Endow, S. A. The Drosophila ncd microtubule motor protein is spindle-associated in meiotic and mitotic cells. J. Cell Sci. 103, 1013–1020 (1992).

8. Walczak, C. E., Verma, S. & Mitchison, T. J. XCTK2: a kinesin-related protein that promotes mitotic spindle assembly in Xenopus laevis egg extracts. J. Cell Biol. 136, 859–870 (1997).

9. Sharp, D. J., Yu, K. R., Sisson, J. C., Sullivan, W. & Scholey, J. M. Antagonistic microtubule-sliding motors position mitotic centrosomes in Drosophila early embryos. Nat. Cell Biol. 1, 51–54 (1999).

10. Goshima, G., Nédélec, F. & Vale, R. D. Mechanisms for focusing mitotic spindle poles by minus end-directed motor proteins. J. Cell Biol. 171, 229–240 (2005).

11. Mountain, V. et al. The kinesin-related protein, HSET, opposes the activity of Eg5 and cross-links microtubules in the mammalian mitotic spindle. J. Cell Biol. 147, 351–366 (1999).

12. Matuliene, J. et al. Function of a minus-end-directed kinesin-like motor protein in mammalian cells. J. Cell Sci. 112, 4041–4050 (1999).

13. Syrovatkina, V. & Tran, P. T. Loss of kinesin-14 results in aneuploidy via kinesin-5-dependent microtubule protrusions leading to chromosome cut. Nat. Commun. 6, 7322 (2015).

14. Kwon, M. et al. Mechanisms to suppress multipolar divisions in cancer cells with extra centrosomes. Genes Dev. 22, 2189–2203 (2008).

15. O’Connell, M. J., Meluh, P. B., Rose, M. D. & Morris, N. R. Suppression of the bimC4 mitotic spindle defect by deletion of klpA, a gene encoding a KAR3-related kinesin-like protein in Aspergillus nidulans. J. Cell Biol. 120, 153–162 (1993).

16. Enos, A. P. & Morris, N. R. Mutation of a gene that encodes a kinesin-like protein blocks nuclear division in A. nidulans. Cell 60, 1019–1027 (1990).

17. Saunders, W. S. & Hoyt, M. A. Kinesin-related proteins required for structural integrity of the mitotic spindle. Cell 70, 451–458 (1992).

18. Olmsted, Z. T., Colliver, A. G., Riehlman, T. D. & Paluh, J. L. Kinesin-14 and kinesin-5 antagonistically regulate microtubule nucleation by γ-TuRC in yeast and human cells. Nat. Commun. 5, 5339 (2014).

19. Paluh, J. L. et al. A mutation in gamma-tubulin alters microtubule dynamics and organization and is synthetically lethal with the kinesin-like protein pkl 1p. Mol. Biol. Cell 11, 1225–1239 (2000).

20. Prigozhina, N. L., Walker, R. A., Oakley, C. E. & Oakley, B. R. Gamma-tubulin and the C-terminal motor domain kinesin-like protein, KLPA, function in the establishment of spindle bipolarity in Aspergillus nidulans. Mol. Biol. Cell 12, 3161–3174 (2001).

21. Wang, B. et al. The Aspergillus nidulans bimC4 mutation provides an excellent tool for identification of kinesin-14 inhibitors. Fungal Genet. Biol. 82, 51–55 (2015).

22. Mieck, C. et al. Non-catalytic motor domains enable processive movement and functional diversification of the kinesin-14 Kar3. Elife 4, 1161 (2015).

23. Jonsson, E., Yamada, M., Vale, R. D. & Goshima, G. Clustering of a kinesin-14 motor enables processive retrograde microtubule-based transport in plants. Nature Plants 1, 1–7 (2015).

24. Fink, G. et al. The mitotic kinesin-14 Ncd drives directional microtubule-microtubule sliding. Nat. Cell Biol. 11, 717–723 (2009).

25. Braun, M., Drummond, D. R., Cross, R. A. & McAinsh, A. D. The kinesin-14 Klp2 organizes microtubules into parallel bundles by an ATP-dependent sorting mechanism. Nat. Cell Biol. 11, 724–730 (2009).

26. Case, R. B., Pierce, D. W., Hom-Booher, N. & Hart, C. L. The directional preference of kinesin motors is specified by an element outside of the motor catalytic domain. Cell 90, 959–66. (1997).

27. Weinger, J. S., Qiu, M., Yang, G. & Kapoor, T. M. A nonmotor microtubule binding site in kinesin-5 is required for filament crosslinking and sliding. Curr. Biol. 21, 154–160 (2011).

28. Stumpff, J. et al. A tethering mechanism controls the processivity and kinetochore-microtubule plus-end enrichment of the kinesin-8 Kif18A. Mol. Cell 43, 764–775 (2011).

29. Sablin, E. P. et al. Direction determination in the minus-end-directed kinesin motor ncd. Nature 395, 813–816 (1998).

30. Henningsen, U. & Schliwa, M. Reversal in the direction of movement of a molecular motor. Nature 389, 93–96 (1997).

31. Endow, S. A. & Waligora, K. W. Determinants of kinesin motor polarity. Science 281, 1200–1202 (1998).

32. Roostalu, J. et al. Directional switching of the kinesin Cin8 through motor coupling. Science 332, 94–99 (2011).

33. Walker, R. A., Salmon, E. D. & Endow, S. A. The Drosophila claret segregation protein is a minus-end directed motor molecule. Nature 347, 780–782 (1990).

34. McDonald, H. B., Stewart, R. J. & Goldstein, L. S. The kinesin-like ncd protein of Drosophila is a minus end-directed microtubule motor. Cell 63, 1159–1165 (1990).

35. Walter, W. J., Machens, I., Rafieian, F. & Diez, S. The non-processive rice kinesin-14 OsKCH1 transports actin filaments along microtubules with two distinct velocities. Nature Plants 1, 15111 (2015).

36. Marcus, A. I., Ambrose, J. C., Blickley, L., Hancock, W. O. & Cyr, R. J. Arabidopsis thaliana protein, ATK1, is a minus-end directed kinesin that exhibits non-processive movement. Cell Motil. Cytoskeleton 52, 144–150 (2002).

37. Ambrose, J. C., Li, W., Marcus, A., Ma, H. & Cyr, R. A minus-end-directed kinesin with plus-end tracking protein activity is involved in spindle morphogenesis. Mol. Biol. Cell 16, 1584–1592 (2005).

38. Furuta, K., Edamatsu, M., Maeda, Y. & Toyoshima, Y. Y. Diffusion and directed movement: in vitro motile properties of fission yeast kinesin-14 Pkl1. J. Biol. Chem. 283, 36465–36473 (2008).

39. Endow, S. A. et al. Yeast Kar3 is a minus-end microtubule motor protein that destabilizes microtubules preferentially at the minus ends. EMBO J. 13, 2708–2713 (1994).

40. Hepperla, A. J. et al. Minus-end-directed Kinesin-14 motors align antiparallel microtubules to control metaphase spindle length. Developmental Cell 31, 61–72 (2014).

41. Scheffler, K., Minnes, R., Fraisier, V., Paoletti, A. & Tran, P. T. Microtubule minus end motors kinesin-14 and dynein drive nuclear congression in parallel pathways. J. Cell Biol. 209, 47–58 (2015).

42. Sproul, L. R., Anderson, D. J., Mackey, A. T., Saunders, W. S. & Gilbert, S. P. Cik1 targets the minus-end kinesin depolymerase kar3 to microtubule plus ends. Curr. Biol. 15, 1420–1427 (2005).

43. Yukawa, M., Ikebe, C. & Toda, T. The Msd1-Wdr8-Pkl1 complex anchors microtubule minus ends to fission yeast spindle pole bodies. J. Cell Biol. 209, 549–562 (2015).

44. Shen, K.-F. & Osmani, S. A. Regulation of mitosis by the NIMA kinase involves TINA and its newly discovered partner, An-WDR8, at spindle pole bodies. Mol. Biol. Cell 24, 3842–3856 (2013).

45. Gerson-Gurwitz, A. et al. Directionality of individual kinesin-5 Cin8 motors is modulated by loop 8, ionic strength and microtubule geometry. EMBO J. 30, 4942–4954 (2011).

46. Fridman, V. et al. Kinesin-5 Kip1 is a bi-directional motor that stabilizes microtubules and tracks their plus-ends in vivo. J. Cell Sci. 126, 4147–4159 (2013).

47. Edamatsu, M. Bidirectional motility of the fission yeast kinesin-5, Cut7. Biochem. Biophys. Res. Commun. 446, 231–234 (2014).

48. Hyman, A. A. Preparation of marked microtubules for the assay of the polarity of microtubule-based motors by fluorescence. J. Cell Sci. Suppl. 14, 125–127 (1991).

